# Males deploy multifaceted strategies and hijack longevity pathways to induce premature demise of the opposite sex

**DOI:** 10.1101/2020.06.30.181008

**Authors:** Lauren N. Booth, Cheng Shi, Cindy Tantilert, Robin W. Yeo, Katja Hebestreit, Cecilia N. Hollenhorst, Coleen T. Murphy, Anne Brunet

## Abstract

Interactions between the sexes negatively impact health in many species, including mammals^1–9^. In mice, sexual interactions induce weight gain and shorten lifespan in females, independent of fertilization^6,9^. In *Caenorhabditis*, males shorten the lifespan of the opposite sex (females or hermaphrodites)^1–3,8^. However, the mechanisms underlying the negative influence of males on lifespan – and their overlap with known longevity pathways – are still largely unknown. Here, we use transcriptomic profiling and targeted screens to systematically uncover new conserved genes involved in male-induced demise. Interestingly, deficiency of these genes individually, and especially in combination, induces strong protection, highlighting the benefit of combining interventions to extend lifespan. Some genes (e.g. *acbp-3, col-43*) only extend hermaphrodite lifespan when knocked-down in the presence of males, suggesting specific protective mechanisms against male-induced demise. However, we also uncover two previously unknown longevity genes (*sri-40* and *delm-2*) that, when knocked-down, extend hermaphrodite lifespan both with and without males, which points to new broad mechanisms of resistance. In sharp contrast, many classical long-lived mutants are actually short-lived in the presence of males, suggesting that males hijack and suppress known longevity pathways. This systematic analysis reveals striking differences in longevity in single sex versus mixed sex environments and uncovers the elaborate network of functional regulation elicited by sexual interactions, which could extend to other species.

## Main

Sexual interactions influence organismal health independently of reproduction in nematodes, flies, and mammals^1–9^. However, the impact of sexual interactions on health is still largely uncharacterized, in large part because most experiments are conducted in single-sex environments. Because of its short lifespan, *C. elegans* is an ideal model organism to systematically examine how sexual interactions affect longevity. Sexual interactions with males shorten the lifespan of the opposite sex (females or hermaphrodites) in *Caenorhabditis*^1–3,8^, and this phenomenon has been shown to involve components from both sexes. Indeed, males promote the premature death of hermaphrodites using male sperm and seminal fluid during mating^2^ as well as male pheromones and secreted compounds^1,10^ (especially when large numbers of males are present^11^). In hermaphrodites, several molecular and cellular pathways have been found to mediate aspects of male-induced demise, including transcription factors (e.g. FOXO/DAF-16 and TFEB/HLH-30)^2,12^, chromatin regulators (e.g. KDM6A/UTX-1)^1^, insulin ligands (e.g. INS-11 and INS-7)^1,2,12^, and even self-sperm itself^12,13^. However, a systematic investigation of the pathways driving the negative impact of males on lifespan, and how they overlap with known longevity pathways, is still missing.

To determine in a systematic manner the impact of sexual interactions with males on the opposite sex in *C. elegans*, we performed RNA sequencing (RNA-seq) of young (day three of life) and middle-aged (day seven of life) hermaphrodites cultured either in the presence or absence of males (Fig. 1a, see Materials and Methods). The males were present for a brief exposure (one day) or for a long exposure (five consecutive days) (Fig. 1a). To eliminate possible effects of sexual interactions during development^10,14,15^ and avoid confounds from embryo transcripts, we initiated exposure to males at the onset of adulthood in sterile (*glp-1[e2144]^16^*) hermaphrodites (Fig. 1a). Male-induced demise occurs robustly and reproducibly under these conditions (Fig. 1b)^1,2^.

**Figure 1:**
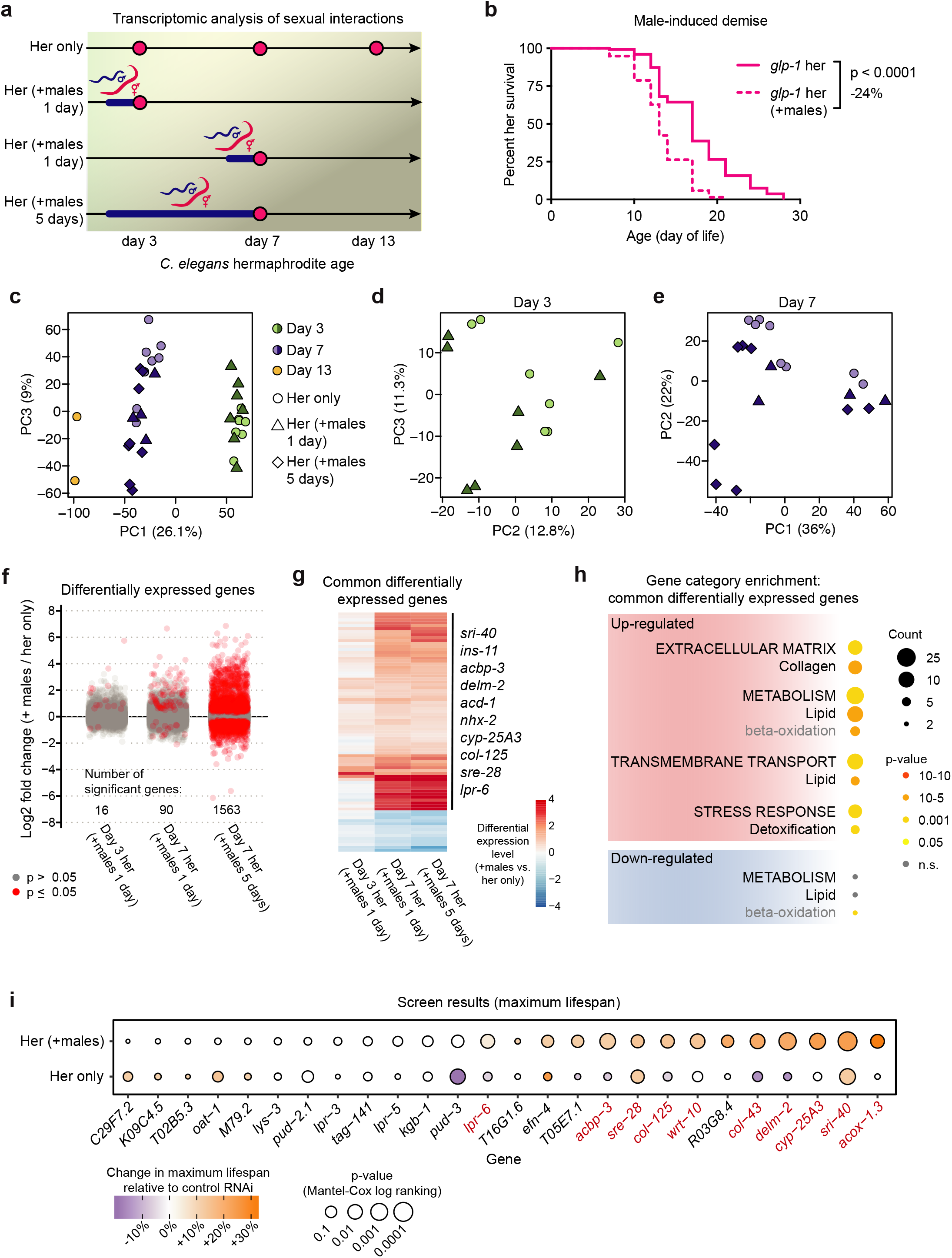
Sexual interactions induce premature death and specific transcriptional changes in *C. elegans* hermaphrodites. (a) Scheme describing the ages of the *glp-1(e2144)* hermaphrodites (Her) used in the RNA-seq experiments and the length of sexual interactions with males. The ages (day of life) of the hermaphrodites at the time of sample collection are indicated with a pink dot and the period of time (one day vs. five days) with males is indicated by a blue line. (b) The presence of males starting at adulthood shortens the lifespan of sterile *glp-1(e2144) C. elegans* hermaphrodites (*p* < 0.0001, −24% change in median lifespan). Lifespan data are plotted as Kaplan-Meier survival curves, the *p*-value was calculated using Mantel-Cox log-ranking, and 79-154 hermaphrodites were tested. A complete list of all lifespan assay results is in Supplementary Table 1. (c-e) Principal Component Analysis (PCA) of the normalized read counts from the hermaphrodite transcriptomes after removal of the male-enriched genes (see Materials and Methods and Extended Data Fig. 1a, b). In panel c, the data from all hermaphrodite age groups are used and in panels d and e the read counts from only day three or day seven were normalized and analyzed. (f) The male-induced hermaphrodite gene expression changes following one day or five days of interaction between the sexes. After filtering the data for male-enriched genes, the log_2_(fold change) for all detected genes is displayed and genes that are significantly differentially expressed (p ≤ 0.05) in the presence of males are shown in red. Complete differentially expression analysis results are in Supplementary Table 2. (g) A heatmap of the genes that are significantly differentially expressed in hermaphrodites in response to the presence of males for both one day (day 3 or 7 hermaphrodites) and five days. The data are displayed as log_2_(fold change). (h) Selected, enriched gene categories from the differentially expressed genes that are common between hermaphrodites that interact with males for one and five days. Gene annotations are nested with the broadest categories listed in all capital letters and the middle categories listed with the first letter capitalized, and the most specific categories in grey. Complete gene set enrichment analysis results are in Supplementary Table 3. (i) The results of our RNAi-based screen for functionally important hermaphrodite genes in male-induced demise. The colors of the circles indicate the change in maximal lifespan relative to control, empty vector RNAi. The sizes of the circles indicate the p-values calculated using Mantel-Cox log ranking. Genes are ordered from smallest to largest change in maximal lifespan in the presence of males, using *p*-value to break ties. The screen hits are indicated with red gene name labels. A complete list of all lifespan data is in Supplemental Table 1.

Principal component analysis separated the transcriptomes of the samples based on age (Fig. 1c, Extended Data Fig. 1a) and exposure to males, especially after longer exposure (Fig. 1d-e). Prolonged exposure to males resulted in a marked increase in the degree and number of gene expression changes compared to the brief exposure to males (Fig. 1f). The genes that were induced in hermaphrodites following a brief (one day) or long (five days) interaction with males partially overlap (e.g. *ins-11*, *sri-40*, *acbp-3*) (Fig. 1g). Importantly, many of the gene expression changes that we observed in sterile, *glp-1* hermaphrodites following sexual interactions with males were also observed in wild-type and fertile hermaphrodites (Extended Data Fig. 1c-e)^13^. Genes up-regulated in response to males were enriched for lipid metabolism and transport, collagens, and stress response (detoxification) gene categories (Fig. 1h, Extended Data Fig. 1e). Thus, males induce transcriptional changes in hermaphrodites, with a greater number and magnitude of changes in response to long interactions.

As both brief and prolonged exposure to males shortens hermaphrodite lifespan^1,2^ (Supplementary Table 1), genes that are induced in both conditions might be more functionally relevant to elicit demise and we focused on these for the remainder of the study. To understand the role of these genes in the male-induced demise of hermaphrodites, we performed a targeted, RNAi-based screen (Fig. 1i and Extended Data Fig. 2). Using wild-type, fertile hermaphrodites, we individually knocked-down male-induced hermaphrodite genes identified through RNA-seq and measured hermaphrodite lifespan in the presence and absence of males. Our screen newly identified ten genes (out of 26 genes tested) that, when knocked-down, partially protected hermaphrodites from male-induced demise (*acbp-3, acox-1.3, col-43,col-125, cyp-25A3, delm-2, lpr-6, sre-28, sri-40*, and *wrt-10*) (Fig. 1i and Extended Data Fig. 2, Supplementary Table 1, see also Fig. 2-4 for further analysis of these genes). Together, these results show striking transcriptional effects of sexual interactions on the opposite sex and identify a set of genes that are functionally important regulators of lifespan in response to sexual interactions.

**Figure 2:**
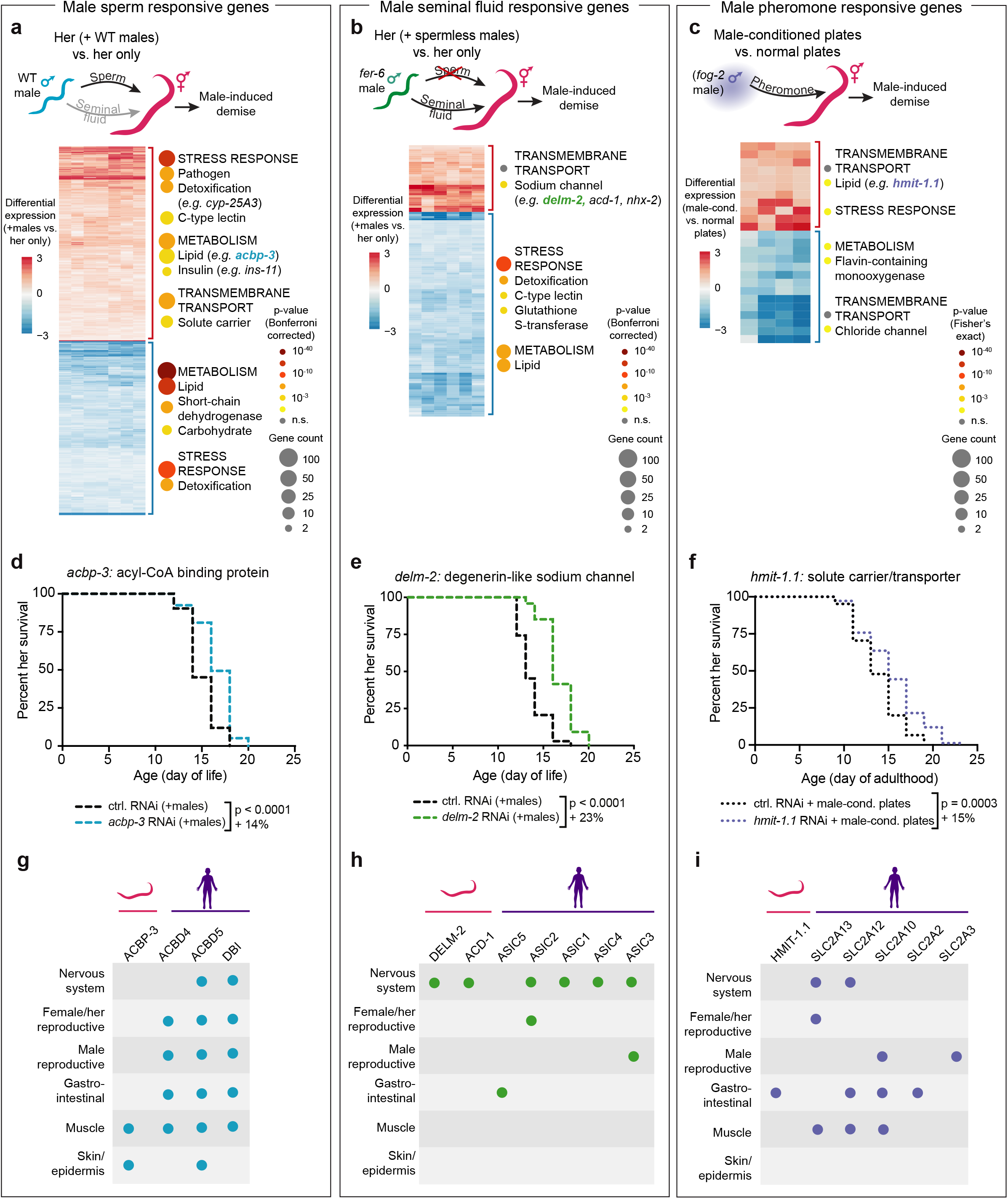
Identification of male sperm, seminal fluid, and pheromone induced genes. (a-c) Heatmaps and gene set enrichment of the microarray results. Heatmap data are displayed as log_2_(fold change). Selected, enriched gene categories for the upregulated (red bracket) and downregulated (blue bracket) genes are shown to the right of each heatmap. Gene category annotations are nested with the broadest categories listed in all capital letters and categories within the broadest categories listed below, with the first letter capitalized. The number of differentially expressed genes in indicated by the size of the circles and enrichment significance by color. The complete results from the microarrays and the gene set enrichment can be found in Supplementary Tables 6 to 9. (a) The male sperm-regulated hermaphrodite *glp-1[e2141]* genes that were identified by microarray as significantly changing in response to mating with wild-type males (that transfer sperm and seminal fluid) but not changing expression in response to mating with males that do not make sperm (*fer-6[hc6]*). The WT male mated hermaphrodite microarray results are shown. (b) A heatmap of the putative male seminal fluid-regulated hermaphrodite genes that were identified by microarray as significantly changing both in response to mating with WT males that transfer sperm and seminal fluid and also with males that transfer seminal fluid but do not make sperm (*fer-6[hc6]*). The gene expression results from *fer-6* male mated hermaphrodites are shown. (c) A heatmap of the hermaphrodite genes that are significantly differentially expressed following exposure to male pheromones (from male-conditioned plates) compared to normal plates. Note that the *p*-values displayed in panel c are from the Fisher’s Exact Test whereas those in panel a and b are the more stringent Bonferroni corrected *p*-values. (d) RNAi knock-down of the sperm-induced gene *acbp-3* extends the lifespan of hermaphrodites in the presence of males (blue dashed line) compared to empty vector control (black dashed line) in the presence of males (*p* < 0.0001). (e) RNAi knock-down of the seminal fluid-induced gene *delm-2* (green dashed line) extends the lifespan of hermaphrodites experiencing a sexual interaction with males (*p* < 0.0001). We note that the *delm-2* targeting RNAi construct is highly similar to paralogs of *delm-2—acd-1* and *delm-1* (Extended Data Fig. 5 and 6). (f) RNAi knock-down of the male-conditioned plate induced gene *hmit-1.1* (purple dashed line) extends the lifespan of hermaphrodites when they are cultured on male-conditioned plates (*p* = 0.0003). Lifespan data for panels d-f are displayed as Kaplan-Meier survival curves and *p*-values calculated using Mantel-Cox log-ranking. Percent change in median lifespan is shown for each lifespan compared to control. 107-124 hermaphrodites were used in each condition. A complete list of all lifespan assay results is in Supplemental Table 1. (g-i) The tissues that express the male-induced genes *acbp-3* (panel g), *delm-2* (panel h), and *hmit-1.1* (panel i) and their human orthologs. The human orthologs shown are the top 5 BLASTp hits. Tissue-specific protein expression is indicated for each ortholog in *C. elegans* (www.wormbase.org, version WS275) and humans (www.proteinatlas.com^46,47^).

We wondered whether the genes we identified as functional mediators of the negative impact of males on lifespan are induced differently by the various male components known to contribute to premature demise (male sperm, seminal fluid, and pheromones^1,2,11^), as this could lead to the development of combined interventions against male-induced demise. To this end, we measured gene expression in young (day 4-5 of life) hermaphrodites following one day of mating with both wild-type males and sperm-less (*fer-6[hc6]^17^)* males (Fig. 2a,b and Extended Data Fig. 3a-d) (thereby distinguishing male sperm from seminal fluid), or on male-conditioned plates for five days (isolating male pheromones) (Fig. 2c). Interestingly, each male component elicited a distinct transcriptional response from hermaphrodites (Extended Data Fig. 3e), consistent with the different physiological changes (*e.g.* fat loss and body shrinking) observed in response to these male components^2,11^.

The subset of genes induced by male sperm (i.e. expressed in response to mating with wild-type but not sperm-deficient males) were enriched for metabolism genes, and included an acyl-CoA binding protein gene (*ACBD4,5/acbp-3)* and an insulin peptide involved in innate immunity (*ins-11)*^18^ (Fig. 2a). RNAi knock-down of the male sperm-dependent genes *acbp-3* (Fig. 2d) and *ins-11*^1^ (as well as other male sperm induced genes) each protected hermaphrodites from male-induced demise (increasing median lifespan by 14-25%, see Supplementary Table 1). Thus, *acbp-3* is a novel gene involved in premature demise via male sperm.

The subset of genes that were likely induced by male seminal fluid (i.e. expressed both in response to mating with wild-type males as well as *fer-6(hc6)* males that cannot make sperm but still transfer seminal fluid^17^) were enriched for sodium channel proteins and include the degenerin-like ion channel genes *ASIC1-5/delm-2* and *acd-1*^19,20^ (Fig. 2b). RNAi knock-down of *delm-2* (which likely also targets *delm-1* and *acd-1* paralogs, Extended Data Fig. 4) and *acd-1* (and other seminal fluid induced genes) each partially protected hermaphrodites from male-induced demise (8-29% increase in median lifespan, see Supplementary Table 1) (Fig. 2e and Fig. 4d). Thus, *delm-2* and *acd-1* (and possibly their paralog *delm-1*), as well as *sri-40* play functional roles in premature hermaphrodite demise via male seminal fluid.

Male-secreted compounds, including pheromones, can shorten hermaphrodite lifespan in the absence of mating^1,10,11^. To identify genes involved in the response of male-secreted compounds (including pheromones), we exposed young, wild-type hermaphrodites to male-conditioned plates and performed a microarray on the hermaphrodites following five days of exposure (Fig. 2c). Genes induced by male-conditioned plates include a solute carrier/transporter gene (*hmit-1.1*). Interestingly, RNAi knock-down of *hmit-1.1* (and other male-conditioned plate induced genes) protected hermaphrodites from the lifespan shortening effect of male-conditioned plates (up to a 17% increase in median lifespan, see Supplementary Table 1) (Fig. 2f and Extended Data Fig. 5). Thus, *hmit-1.1* and *C33G8.3* are novel regulators of hermaphrodite demise induced by male-secreted compounds, including male pheromones.

Interestingly, many of the genes induced by the different male components are conserved between *C. elegans* and humans. Some of the human genes are expressed in the same tissues as their *C. elegans* orthologs (Fig. 2g-i and Extended Data Fig. 6), where they could initiate a conserved response to sexual interactions. Collectively, these findings show that male sperm, seminal fluid, and pheromones induce the expression of different subsets of novel and conserved genes that functionally contribute to premature demise in the opposite sex.

The observation that different genetic pathways mediate the demise induced by male sperm, seminal fluid, and pheromones raises the possibility that modulating these genes in combination could be more effective to counter male-induced demise (Fig. 3a). To test this, we performed lifespan assays in which we combined losses of gene function. We assessed the role of the gene *delm-2* (induced by male seminal fluid) in combination with *acbp-3* (induced by male sperm) (Fig. 3b). While knock-down of each gene individually resulted in a partial protection from male-induced demise, knock-down of *delm-2* and *acbp-3* together resulted in additive protection against male-induced demise (11-31% increase in median lifespan compared to a single gene knock-down, see Supplementary Table 1) (Fig. 3b). Interestingly, knock-down of *delm-2* and *acbp-3* in combination fully protected hermaphrodites from the lifespan shortening impact of sexual interactions with males (43-50% increase in median lifespan compared to control RNAi, see Supplementary Table 1) (Fig. 3c), and these hermaphrodites lived approximately the same lifespan as hermaphrodites in the absence of males (Fig. 3c, Supplementary Table 1). Thus, a combined loss of function in the male sperm and male seminal fluid pathways improves protection from male-induced demise. Analysis of the regulatory regions of the genes induced by parallel male strategies revealed key transcription factor binding sites (Fig. 3d), with some of them being shared among different pathways (e.g. PQM-1^12^, PPARD/NHR-28, and conserved PBX3,4/CEH-60) (Fig. 3d, Extended Data Fig. 7). Together, these results suggest that targeting the different strategies induced by males in combination (using multiple genes and perhaps common transcription factors) is more effective to counter male-induced demise than targeting a single strategy.

**Figure 3:**
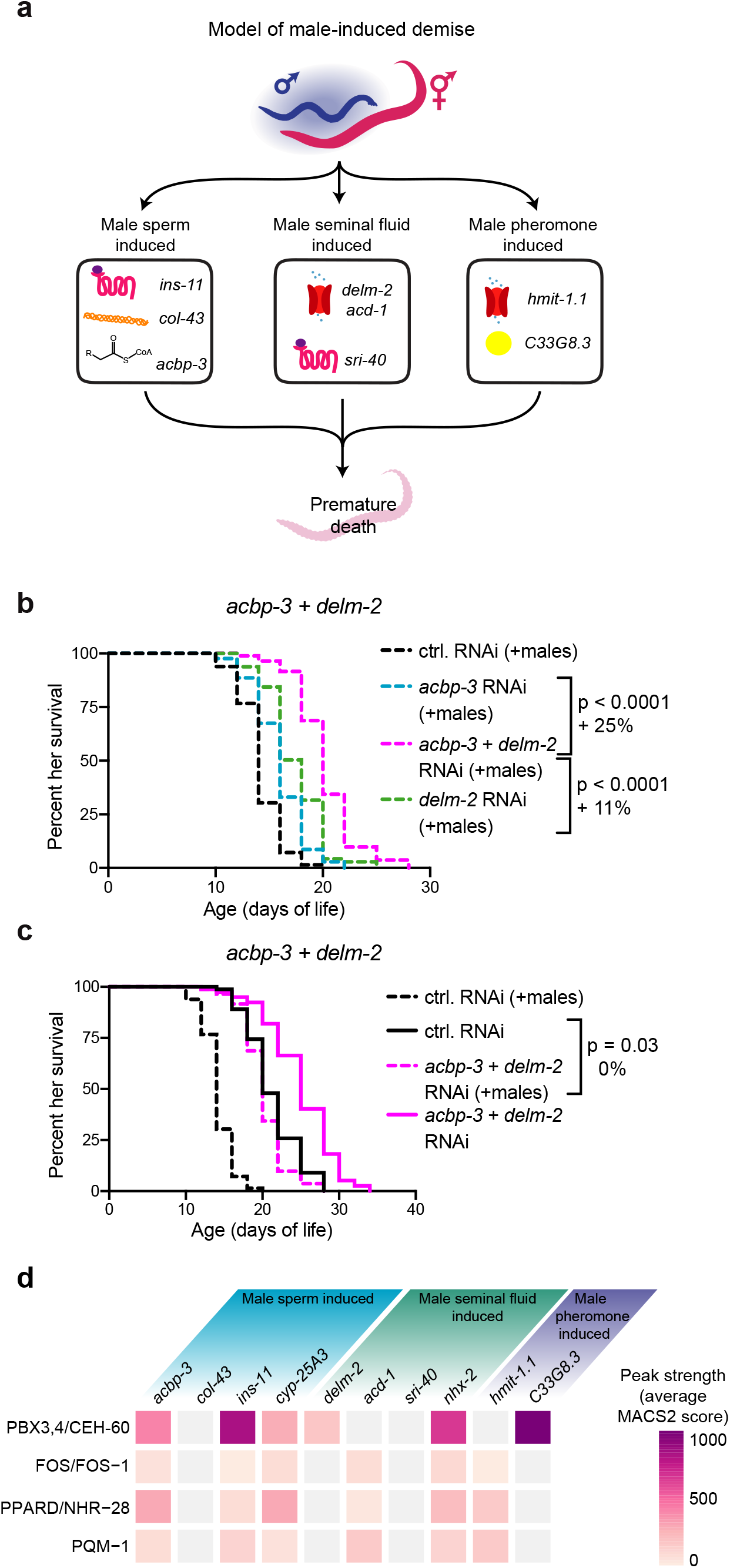
Strategies to target multiple male-induced demise pathways. (a) A summary of the transcriptomics and male-induced demise lifespan data showing that male sperm, seminal fluid, and pheromones each induce a different set of functionally important male-induced demise gene in hermaphrodites. (b) Knock-down of *acbp-3* or *delm-2* individually by RNAi partially protected hermaphrodites from male-induced demise (*acbp-3: p* < 0.0001, 14% increase in median lifespan and *delm-2: p* < 0.0001, 29% increase in median lifespan compared to control RNAi). Knock-down of *acbp-3* and *delm-2* simultaneously by double RNAi (dashed pink line) protected hermaphrodites from male-induced demise to a greater extent than knock-down of either *acbp-3* or *delm-2* alone (*p* < 0.0001, 25% and 11% increase in median lifespan, respectively). (c) Knock-down of *acbp-3* and *delm-2* simultaneously extended hermaphrodite lifespan. In the presence of males (the dashed lines), loss of *acbp-3* and *delm-2* increased median lifespan by 43% (*p* < 0.0001). This resulted in a lifespan that was comparable to control hermaphrodites in a single-sex setting (black, solid line vs. dashed, pink line: *p =* 0.03, no change in median lifespan). In the absence of males, loss of *acbp-3* and *delm-2* increased median lifespan by 25% (*p* < 0.0001) compared to control RNAi (solid, black line). Lifespan data are plotted as Kaplan-Meier survival curves, the *p*-value was calculated using Mantel-Cox log-ranking, and percent change in median lifespan is shown. 105-114 hermaphrodites were tested. A complete list of all lifespan data is in Supplementary Table 1. (d) The average MACS2 score of transcription factor ChIP-seq binding peaks within 5kb +/− of the transcription start site of male-induced functional important genes. A complete list of the results of the transcription factor enrichment analysis can be found in Supplementary Table 10.

We next wondered whether genes that mediate male-induced demise are specific to the presence of males or whether they have a more general effect on lifespan, and how this compares to traditional longevity mutants (Fig. 4a). Several of the male-induced genes we identified specifically extended lifespan when knocked-down only in the presence of males (e.g. *acbp-3, col-43*), and did not confer a longer hermaphrodite lifespan in the absence of males (Fig. 4a-e). These genes represent pathways that are specifically involved in regulating longevity in a mixed sex environment.

**Figure 4:**
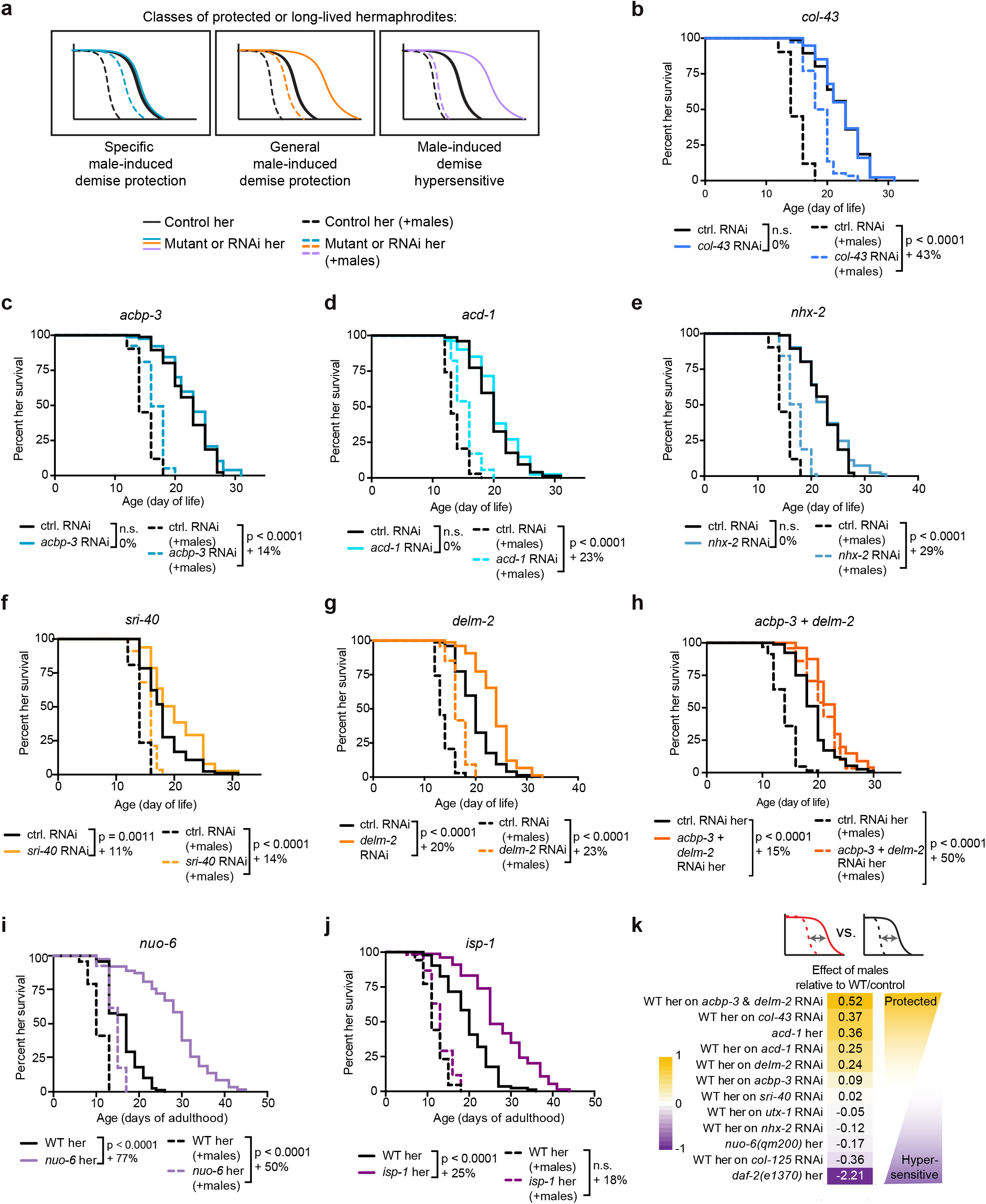
Male-induced demise is mediated by specific genes, general health genes, and hijacking of conserved longevity pathways. (a) A scheme describing the different effects of lifespan extending knock-down and mutations in the presence and absence of males. (b-e) Knock-down of the collagen gene *col-43* (b), the acyl-CoA binding protein gene *acbp-3* (c), the ion channel *acd-1* (d), or the transporter *nhx-2* (e) by RNAi partially protected hermaphrodites from male induced demise (dashed lines) but did not detectably extend lifespan in the absence of males (solid lines). We note that the male-induced demise lifespans in panel c (dashed lines) are also displayed in Fig. 2d and are displayed again here for comparison with hermaphrodite lifespan in the absence of males. (f-g) Knock-down of the serpentine receptor *sri-40* (f) or the sodium channel *delm-2* (g) extended hermaphrodite lifespan in both the presence (dashed lines) and absence (solid lines) of males. We note that the male-induced demise lifespans in panel g (dashed lines) are also displayed in Fig. 2e and are displayed again here for comparison with the lifespans in the absence of males. (h) Knock-down of both *delm-2* and *acbp-3* simultaneously extended hermaphrodite lifespan in both the absence of males (solid lines) and in the presence of males (dashed lines). (i-j) Mutations in the mitochondria electron transport chain genes *nuo-6(qm200)* and *isp-1(qm150)* extended lifespan in a hermaphrodites only setting (solid lines). In the presence of males, *nuo-6* hermaphrodites lived longer than wild-type (panel e, dotted lines), but this extension was blunted compared to the extension of lifespan in the absence of males (see panel k). In the presence of males, *isp-1* mutant hermaphrodites did not live longer than wild-type (panel j, dashed lines). All lifespan data are plotted as Kaplan-Meier survival curves, the *p*-values were calculated using Mantel-Cox log-ranking, and percent changes in median lifespan are shown for each comparison. 55-116 hermaphrodites were tested. A complete list of all lifespan data is in Supplementary Table 1. (k) A summary of the percent loss of lifespan in the presence of males in the mutant or RNAi knock-down compared to wild-type. The effects of males for each genetic manipulation relative to wild-type was calculated as 1 – (male-induced change in median lifespan of hermaphrodites with mutation or RNAi / male-induced change in median lifespan of control). Positive values indicate that the genetic manipulation resulted in greater protection from male-induced demise whereas negative values indicate hypersensitivity to male-induced demise. The *daf-2(e1370)* results were published previously^1^. All other lifespan results are from this manuscript (see Supplementary Table 1 and 11).

Other genes lead to lifespan extension when knocked-down both in absence and presence of males. We previously found that *utx-1* RNAi knock-down extends hermaphrodite lifespan in the absence of males^21,22^ and protects hermaphrodites from male-induced demise^1^. Here, we identify three additional genetic interventions that protected from male-induced demise and promoted longevity in the absence of males: RNAi knock-down of *delm-2* or *sri-40* individually (Fig. 4a, f, g) and knock-down of *delm-2* and *acbp-3* in combination (Fig. 4h). These are genes that can regulate longevity in both a single and mixed sex environment.

In sharp contrast, traditional long-lived mutant hermaphrodites (*daf-2[e1370]^23,24^, glp-1[e2144]^25^*) are hypersensitive to male-induced demise^1,2^. Similar to *daf-2* mutants, we find that long-lived mitochondria mutants (e.g. *isp-1*^26^, *nuo-6*^27^) are also hypersensitive to the presence of males, as their lifespan extension is smaller (or undetectable) in the presence of males compared to the absence of males (Fig. 4a, 4i, j). This may be due to males “hijacking” traditional longevity genes and pathways to repress them, possibly through inhibition of the FOXO/DAF-16 transcription factor^2,23–25,28^. These genes represent pathways that are strongly involved in regulating lifespan in a single sex environment, but not in a mixed sex environment.

Together, these results show that males impinge on the health of the opposite sex in different ways — highly specific pathways (*e.g. acbp-3*), more general health pathways (*e.g. delm-2*), and the “hijacking” of conserved longevity pathways (Fig. 4k and Extended Data Fig. 8). These findings also reveal that the context in which an individual lives (for example, with or without mates) can drastically impact the effect of a longevity mutation.

Here, we have identified genes that are not only regulated by but are also functionally important for the response of hermaphrodites to sexual interactions (male-induced demise). By investigating these genes through different “lenses” we have found that they fall into different functional categories both in terms of the male signal to which they respond (male sperm, seminal fluid, or pheromones) and the specificity of their regulation of lifespan (specific to male-induced demise versus broad longevity regulation). Many of these genes are conserved in mammals, and they could also play a role in lifespan and health in other species, including humans.

The placement of the male-induced demise genes in different pathways allowed us to predict which combination of strategies could lead to greater protection from male-induced demise—knock-down of independent male-induced demise pathways in combination has an additive effect towards protecting hermaphrodites against males. In fact, knock-down of *delm-2* and *acbp-3* in combination extended the lifespan of hermaphrodites in the presence of males to the lifespan of hermaphrodites in the absence of males. This is important because it could help identify additional combinations of strategies to more robustly extend lifespan.

Our analysis identifies genes whose knock-down can extend lifespan both in the absence and presence of males (*e.g. sri-40* and *delm-2*) and that have been missed by previous genetic screens. Thus, elucidating the detrimental effects of sexual interactions on health can also identify pathways that regulate health more generally. These newly-identified regulators of longevity have promise to protect individuals from a range of different stressors, unlike the specific regulators of male-induced demise we have identified (*e.g. acbp-3* and *col-43*). In contrast, several classical longevity mutants are, in fact, short-lived (hypersensitive) in the presence of males. These results suggest that males hijack components of traditional longevity pathways and that long-lived mutants may have an Achilles’ heel – their susceptibility to the opposite sex. Sexual interactions may therefore represent a particularly potent biological force distinct from other types of stresses. Our study also reveals that targeting genes in a mixed vs. a single sex environment can have different outcomes on lifespan, which has important implications for health.

## Acknowledgements

We thank Anne Villeneuve, Miriam Goodman, and Laura Bianchi for helpful discussions. Thank you to all the members of the Brunet lab, particularly Jingxun Chen, Salah Mahmoudi, Jason Miklas, and Katharina Papsdorf for helpful feedback and discussion. Thank you to the members of the Murphy lab for valuable feedback. The Stanford Functional Genomics Facility performed the sequencing of the RNA-seq libraries. We thank the *Caenorhabditis* Genetics Center (funded by NIH grant P40 OD010440), Miriam Goodman, Laura Bianchi, and Man-Wah Tan for providing *C. elegans* strains used in this study and WormBase. This work was funded by R01AG054201 (AB), HHMI-Simons Faculty Scholars Program (CTM), Glenn Foundation for Medical Research (CTM), Genentech Foundation Predoctoral Fellowship (RWY), the Helen Hay Whitney Foundation (LNB), and NIH K99 AG051738 (LNB).

## Author Contributions

This study was designed by LNB, CS, CTM, and AB. LNB performed the RNA-seq experiments and data were analyzed by LNB and KH. Microarrays were performed by CS and analyzed by CS, LNB, and RWY. All LNB and KH-written code was independently checked by RWY. Lifespan assays were performed and analyzed by LNB, CS, and CT (see Supplementary Table 1 for the assays that were performed by each researcher). CNH performed assay validation. The figures were prepared by LNB with feedback from AB, CTM, and CS. The original manuscript draft was written by LNB with advice and editing from AB, CTM, and CS.

## Competing Interests

The authors declare no competing interests.

## Data Availability

All RNA-seq reads are available on NCBI Sequence Read Archive (PRJNA642294). The results from the microarrays are available at http://puma.princeton.edu (see Materials and Methods). The complete list of all lifespan assays (including statistics and number of animals) is presented in Supplementary Table 1 and the results of our RNA-seq and microarray analyses are presented in Supplementary Tables 2 to 10.

## Code Availability

All code is available on GitHub (https://github.com/brunetlab/Booth-et-al.-2020).

## Materials and Methods

### Worm strains and maintenance

All *C. elegans* wild-type and mutant strains used in this study are listed below. All strains were maintained on Nematode Growth Media (NGM) plates with 50μg/mL streptomycin (Gibco) and a lawn of OP50-1 bacteria (a gift from M.-W. Tan) from stationary phase cultures. Nematodes were grown at 20°C, with the exception of temperature-sensitive mutants (*glp-1[e2144]* and *[e2141]*), which were maintained at 15°C (permissive temperature). When temperature sensitive mutants were used for assays, they were grown at the restrictive temperature (25°C). The genotype of strains was verified by genotyping PCR and Sanger sequencing and the strains were backcrossed three times into our laboratory’s N2 strain (in addition to the backcrossing that was performed when the mutants were initially isolated).

| *C. elegans* Strain | Genotype | Source |
| --- | --- | --- |
| N2 | wild-type | Dr. Man-Wah Tan and CGC |
| CF1903 | *glp-1(e2144) III* | CGC |
| CB4037 | *glp-1(e2141) III* | CGC |
| CB1467 | *him-5(e1467) V* | CGC |
| BA6 | *fer-6(hc6) I* | CGC |
| CB4108 | *fog-2(q71) V* | CGC |
| RB1523 | *delm-2(ok1822) I* | CGC and CDMC |
| RB1177 | *delm-1(ok1226) IV* | CGC and CDMC |
| BLC100 | *delm-1(ok1226) IV*;<br>*delm-2(ok1822) I* | Dr. Laura Bianchi |
| ZB90 | *acd-1(bz90) I* | Dr. Laura Bianchi |
| CB1611 | *mec-4(e1611) X* | Dr. Miriam Goodman |
| MQ1333 | *nuo-6(qm200) I* | CGC |
| MQ887 | *isp-1(qm150) IV* | CGC |
| CB4876 | *clk-1(e2519) III* | CGC |

\*CGC is the *Caenorhabditis* Genetics Center
\*CDMC is the *C. elegans* Deletion Mutant Consortium<sup>29</sup>

### RNA-seq

To better understand how males induce premature hermaphrodite demise, we characterized the transcriptomes of sterile *glp-1(e2144)* hermaphrodites that have a shortened lifespan after interacting with wild-type males for one day or five days to those that never interacted with males.

Individuals were age-synchronized using a brief, 3-4 hour egg lay (see “Lifespan Assays” below) on 10cm NGM plates seeded with OP50-1 bacteria and grown at 25°C during development and adulthood. The day of the egg lay is considered day 0 of life. For the longer (five day) exposure to males, 75 *glp-1(e2144)* hermaphrodites were placed onto a 10cm NGM plates seeded with OP50-1 bacteria and with 75 young (day 3-5 of life) WT males starting on day 2 of life (young adults) until day 7 of life. At day 5 of life, the hermaphrodites were moved to fresh plates and the males were replaced with new, young WT males. For samples in which hermaphrodites interacted with males for a single day, 75 *glp-1(e2144)* hermaphrodites were placed onto a 10cm NGM plates seeded with OP50-1 bacteria and 75 young (day 3-5 of life) WT males were added starting on either day 2 of life (young adults) or on day 6 of life.

Hermaphrodites that interacted with males starting on day 6 of life were maintained on 10cm NGM plates with OP50-1 at a density of 150 hermaphrodites starting from day 2 of life. At the same time, hermaphrodites from the same cohort of age-synchronized *glp-1(e2144)* individuals were also placed on fresh plates without males (150 hermaphrodites per 10cm plate to maintain a similar density). After one or five days, 75 hermaphrodites from each condition (individuals that either interacted with males or never interacted with males) were collected by removing either 75 hermaphrodites (for the “no males” condition) or the 75 males from the plates. For each sample, these remaining 75 hermaphrodites were immediately washed three times with ice cold M9 buffer (22mM KH_2_PO_4_, 42mM Na_2_PO_4_, 86mM NaCl, and 1mM MgSO_4_) and the worm pellets were flash frozen in liquid nitrogen. In parallel with the RNA-seq sample collection, *glp-1(e2144)* hermaphrodites and WT males form the same batches of age-synchronization as their corresponding RNA-seq samples were used to measure hermaphrodite lifespan as described in the section “Lifespan Assays” (see Supplementary Table 1).

To determine the contribution of male transcripts to the *glp-1(e2144)* samples, we also isolated and prepared for RNA-sequencing the population of WT males that were on the RNA-seq replicates E-H plates with the day 6-7 of life, single day sexual interaction condition. These young male worms (not synchronized) were washed with ice cold M9 buffer and flash frozen as described above.

RNA was extracted from the flash frozen worm pellets (approximately 75 whole worms per sample) with 500μL Trizol and 200μL chloroform followed by 250μL phenol and 200μL chloroform extractions and finally, an isopropanol precipitation. Remaining DNA was degraded with DNaseI (Promega) and the RNA cleaned with a sodium acetate and ethanol precipitation. RNA quality was measured using Nanodrop spectrophotometry and the Agilent BioAnalyzer Total RNA Nano chip and kit. mRNA enriched cDNA was prepared using 10ng of total RNA (quantified by Nanodrop spectrophotometry) and the Takara SMART-seq v4 Ultra Low Input RNA kit, with 8 rounds of amplification. Paired-end libraries were made using the Nextera XT DNA library prep kit (Illumina) with 1ng of cDNA (quantified using the Qubit dsDNA High Sensitivity reagents, Invitrogen) and barcoded using the Nextera XT Index Kit v2 (Illumina). Libraries were purified with 30μL AMPure XP beads (Beckman Coulter) as directed in the Nextera XT kit. Library quality and quantity were assessed using the Agilent Bioanalyxer High Sensitivity DNA Assay. The libraries for biological replicates A-D were prepared together and pooled and sequenced on a single Illumina NextSeq run and the libraries for biological replicates E-H were prepared together and pooled and sequenced on two Illumina NextSeq runs. Paired-end, 75 base pair sequencing was performed. All RNA-seq reads are publicly available through NCBI Sequence Read Archive (BioProject PRJNA642294).

### RNA-seq analysis

RNA-seq reads were aligned to the WBcel235 genome and gene read counts were calculated using STAR (version 2.5.4a)^30^. Low-coverage genes that had less than one read count per million mapped reads in less than three samples were filtered out. After filtering of low-coverage genes, 16,706 genes remained (prior to removal of male-enriched genes, see below) with read coverage per sample ranging from 3,650,980 for the lowest quantile and 31,137,429 for the highest quantile. Data was normalized with a variance-stabilizing transformation (DESeq2 version 1.10.1)^31^ prior to Principal Component Analysis (PCA) in R (version 3.2.4 and Biobase version 2.30.0 and 2.42.0^32^). PCA was carried out using the R method (prcomp). Differential expression was calculated using DESeq2 (version 1.10.1)^31^. The results from DESeq2 can be found in Supplementary Tables 2 and 4. Heatmaps were generated in R using normalized read counts (variance-stabilizing transformation).

Hermaphrodites that have interacted with males receive male sperm as a result of mating and male-sperm derived transcripts were detected in our data (Extended Data Fig. 1). To focus on the effect of sexual interactions of the hermaphrodites, we developed a list of male-enriched genes that were excluded from future analysis. Briefly, we used DEseq2 to calculate differential expression between the WT males and the *glp-1(e2144)* hermaphrodites that never experienced a sexual interaction (day 3 and 7 of life were combined for this analysis). Genes that were expressed more highly in WT males (log_2_(fold change) > 0, adjusted *p*-value ≤ 0.05) were excluded from the datasets. 5,355 genes met this threshold and were called as male-enriched. This filtered dataset was then carried through our standard RNA-seq pipeline (see above). Both the non-filtered datasets and the datasets in which the male-enriched genes were removed from the analysis are available as supplemental data sets and their analyses are included in Supplementary Table 2 and 4.

All code is publicly available online (https://github.com/brunetlab/Booth-et-al.-2020).

### Gene set enrichment

WormCat^33^ was used to determine the gene set enrichment for the RNA-seq and microarray results. Genes were identified as significant by SAM (see below) and DESeq2 (see above). For the RNA-seq, genes were called as differentially expressed if the adjusted *p*-value was less than 0.05. Significantly up- and down-regulated genes were input to WormCat.org using default settings. For gene sets with a very small number of genes (those up- and down-regulated on male-conditioned plates), we did not observe significant enrichment using the multiple-hypothesis corrected *p*-values and instead present the Fisher’s Exact Test for these gene sets. Complete lists of all WormCat gene set enrichment results, including both the Fisher’s Exact Test and Bonferroni corrected *p*-values are presented in Supplementary Tables 3, 5, and 9.

### Lifespan assays

*C. elegans* hermaphrodites used for lifespan assays were age-synchronized with a short (3-4 hour) egg-lay using young (day 3-5 of life), well-fed, adult parents. All worms were grown on NGM plates with streptomycin (50μg/mL) and seeded with OP50-1 bacteria unless RNAi knock-down was performed (see below).

For each assay, worms were scored as dead or alive and transferred to new plates daily during the reproductive period and then every other day. Worms were scored as dead if they did not respond to gentle, repeated prodding with a wire pick (90% Pt, 10% Ir) along different points of their body. Worms were scored as censored if they crawled off the media or died due to bagging (internal hatching) or vulval rupture. Data from these censored worms were included up until the point of censorship (see Supplementary Table 1 for all data).

For conditions in which the effect of sexual interactions was assessed, we used one of three methods, as indicated. For the long-term exposure method (described previously^1,2^), young males (day 1 to 2 of adulthood) were added to the hermaphrodites at the onset of adulthood. For lifespan experiments in which the hermaphrodites were exposed to males for their entire adulthood^1^, males were added in a 1:1 ratio with hermaphrodites and the number of males remained fixed, even as hermaphrodites began to die or censored. Male worms were replaced every other day at the time the hermaphrodites were transferred to new plates. Male stocks were set up approximately every other day for the entirety of the lifespan assay. For the lifespan experiments in which hermaphrodites were exposed to males for only one day^2^, young males were added in a 2:1 male:hermaphrodite ratio. Following 24 hours of exposure, hermaphrodites were moved to new plates and did not encounter a male again throughout their lifespan. For the male-conditioned media lifespan assays, hermaphrodites were transferred on to male-conditioned plates (MCP) from late L4 stage and stayed on MCP for the remainder of their lives. Male-conditioned plates were prepared throughout the course of the lifespan assays: 30 Day 1 of life males (*fog-2(q71)*, essentially WT) were transferred onto each plate (35mm NGM plates). Two days later, the males were removed and hermaphrodites for lifespan assays were immediately transferred onto these male-conditioned plates.

Synchronized worms (hermaphrodites, feminized individuals etc.) were randomly assigned to the “no males” or “+ males” conditions by picking them onto fresh plates in an alternating manner to avoid selection bias. Similarly, the males used for mating with individuals of different genotypes or RNAi treatments were from the same sets of males in each assay and were allocated randomly in an alternating manner. For each single biological replicate, approximately 35 individuals were placed on each of 3-6 plates (each plate represents a technical replicate). The number of individuals per plate and number of technical replicates were chosen based on field standards^34^.

For sterile mutants, slight modifications were made to the methods. The sterile *glp-1(e2144)* mutant and WT control parents were used for an egg lay at the permissive temperature (15°C). Following the egg lay, the individuals used for the assay were kept at 25°C for the remainder of the assay.

Lifespan data were plotted as Kaplan-Meier survival curves in Prism 8 and statistical analyses performed using the logrank (Mantel-Cox) test. The number of animals (n) used for each assay and the number of independent biological replicates (N) can be found in Supplementary Table 1.

### RNAi knock-down

To knock-down expression of specific genes, we fed worms HT115 (*E. coli*) bacteria expressing double strand RNA targeting a specific gene. Worms were cultured on NGM containing ampicillin (100μg/mL, Sigma) and IPTG (0.4mM, Invitrogen). During development, worms were fed HT115 bacteria (grown to stationary phase, RNAi expression induced for 2-4 hours with 0.4mM IPTG, and the bacteria concentrated to 20x) carrying control empty vector (EV). Upon adulthood (day 3 of life), worms were placed onto plates with HT115 bacteria (grown to stationary phase, RNAi expression induced for 2-4 hours with 0.4mM IPTG, and the bacteria concentrated to 20x) carrying the appropriate RNAi clone. RNAi clones in HT115 *E. coli* were isolated from the Ahringer RNAi library^35^ (a gift from A. Fire) or, if unavailable, from the Vidal RNAi library^36^ (Dharmacon). The inserts of the plasmids encoding the RNAi clones used in this study were sequenced to verify their identity. For all lifespan assays, the identity of the RNAi clone was blinded until the lifespan assay was completed.

We note that the RNAi construct that targets *delm-2* shares high sequence similarity to two *delm-2* paralogs—*delm-1* and *acd-1* (Extended Data Fig. 4). Interestingly, *acd-1* expression was also induced by males (Fig. 1g, 2b and Extended Data Fig. 3d) and loss of *acd-1* by RNAi knock-down partially protected hermaphrodites from male-induced demise (Fig. 4c, Extended Data Fig. 4b, and Supplementary Table 1). However, single mutations in these genes are not sufficient to protect hermaphrodites from male-induced demise (Extended Data Fig. 4b-f), suggesting that these channel genes may act redundantly or that the RNAi knock-down results in a greater loss of function either of a single gene or a combination of *delm-2, delm-1*, and *acd-1.* For simplicity, in the manuscript, we refer to this manipulation as *delm-2* knock-down.

To perform double RNAi, we combined equal amounts of bacteria expressing *delm-2* targeting dsRNA and *acbp-3* targeting dsRNA. This was compared to control RNAi bacteria (with empty vector) and to single RNAi knock-down. The single RNAi knock-down for these experiments were diluted 50% using control (empty vector) RNAi expressing bacteria. We note that RNAi knock-down does not result in complete loss of function (i.e. it is not a null). Therefore, a caveat to the interpretation of double RNAi results is that effects on lifespan may be due to intensifying the loss of function in a single pathway rather than targeting two parallel pathways.

### RNAi-based screen

To determine whether the male-induced gene expression changes in hermaphrodites functionally contribute to their premature demise, we performed a targeted RNAi-based screen. The specific genes that we included in our screen were chosen because they were up-regulated in multiple datasets: *glp-1* sterile hermaphrodites for long and short interactions (Fig. 1g) and WT or feminized individuals mated with males for two hours^13^ (Extended Data Fig. 1d). We also included several genes that were highly male-enriched (*lys-3, C29F7.2, T02B5.3*, and *T16G1.6*) and that were not significantly enriched (*kgb-1* and *K09C4.5*) as controls to test whether genes that we filtered from our RNA-seq analysis (see above) were functionally important for male-induced demise. None of these genes significantly extended hermaphrodite lifespan when knocked-down (Supplementary Table 1). As a positive control, we used RNAi knock-down of the male-induced demise gene *utx-1^1^*. RNAi treatments were blinded until the last animals died.

For the screen, we performed lifespan assays using a single plate of approximately 35 N2 (wild-type) hermaphrodites for each “no males” condition and two plates of approximately 18 N2 (wild-type) hermaphrodites and 18 *him-5(e1467)* males for each “+ males” condition. For details on the lifespan assays, see above.

### Microarrays

*glp-1(e2141)* hermaphrodites were mated with wild-type or *fer-6(hc6)* males starting on day 1 of adulthood for 24 hours in a 2:1 male to hermaphrodite ratio. *fer-6(hc6)* males do not transfer sperm but have normal seminal fluid and copulation during mating^3,17^. 200 hermaphrodites were collected on day 2/3 of adulthood. Four biological replicates of wild-type males mated *glp-1* hermaphrodites were collected on day 2 of adulthood, another three replicates were collected on day 3 of adulthood. All six replicates of *fer-6* male mated *glp-1* hermaphrodites were collected on day 3 of adulthood.

Because male pheromone-induced demise depends on an intact germline^11^, male pheromone-induced expression changes might be missed in our RNA-seq and microarray transcriptomic measurements using *glp-1* (germline-less) animals. Therefore, we identified male pheromone-induced expression changes using WT N2 hermaphrodites. Synchronized late L4 N2 hermaphrodites were picked on to 35 mm plates (control and male-conditioned plates [conditioned by 30 young *fog-2* males for two days, see above]). 30 worms per plate—about 180 worms in total—were used for each biological replicate. Hermaphrodites were transferred on to freshly seeded or male-conditioned plates every two days, and collected on day six for RNA extraction. Four biological replicates were performed. Interestingly, male-conditioned plates induced hermaphrodite gene expression changes that were very similar to the transcriptional response of males to male-conditioned plates (up-regulated genes *p*-value = 0.00057 and down-regulated genes *p*-value = 4.96 × 10^−5^, hypergeometric test)^11^.

RNA was extracted by the heat vortexing method. Two-color Agilent microarrays were used for expression analysis. Significantly differentially expressed gene set were identified using SAM^37^. Additional analysis was performed in R and all R code is publicly available online (https://github.com/brunetlab/Booth-et-al.-2020).

Microarray data can be found in Supplementary Tables 6-8 and PUMAdb (http://puma.princeton.edu):

*glp-1* hermaphrodites mated with wild-type males: https://puma.princeton.edu/cgi-bin/exptsets/review.pl?exptset_no=7345
*glp-1* hermaphrodites mated with *fer-6* males: https://puma.princeton.edu/cgi-bin/exptsets/review.pl?exptset_no=7346
N2 hermaphrodites on male-conditioned plates: https://puma.princeton.edu/cgi-bin/exptsets/review.pl?exptset_no=7351

### Comparison of gene expression results

The transcriptomic experiments (Fig. 1 & 2) were performed by different researchers using different experimental setups (e.g. the ratio of males to hermaphrodites) and nematode strains. To compare the RNA-seq and microarray gene expression results, we: calculated Pearson’s correlations of the fold changes for the genes measured in the different experiments (Extended Data Fig. 3a), identified the genes that are common between the microarray and RNA-seq results and performed hypergeometric tests (Extended Data Fig. 3b), and compared gene set enrichments (Extended Data Fig. 3c). These comparisons revealed that the RNA-seq and microarray yielded highly similar results at the whole-genome and specific gene levels, highlighting the robustness of the male-induced demise phenomenon and our data.

### Alignment of *delm-2* RNAi targeting sequence

The *delm-1* and *acd-1* unspliced transcript sequences were downloaded from WormBase (WS275) and aligned to the Ahringer library RNAi construct that targets *delm-2* using Clustal Omega^38^. The pairwise alignments were visualized using JalView^39^.

### Transcription factor binding enrichment

ChIP-Atlas^40^ was used to determine the enrichment of ChIP-seq peaks at the male sperm, seminal fluid, and pheromone regulated genes. These differentially expressed genes were identified by microarrays (see above). The ChIP-seq peak significance threshold was set to 100 (equivalent to a Q-value < 1 × 10^−10^) and ChIP-seq peaks from the groups “TFs and others” and “All cell types” were used. ChIP-seq peaks within 2kb^41^ up- and downstream of the transcription start site were considered for each differentially expressed gene. The presence and strength of transcription factor peaks for select genes (Fig. 3d) was also determined using ChIP-Atlas on default settings. We note that these ChIP-seq experiments were performed in whole worms at various developmental stages and under different conditions. Thus, it is important to keep in mind that these transcription factor-gene regulatory connections may not represent the situation in a given adult tissue undergoing male-induced demise. The complete list of enriched transcription factor binding is in Supplementary Table 10.

### Motif analysis

To calculate *known* and *de novo* motif enrichment within the promoter regions of differentially expressed genes, we used the Homer^42^ function “findMotifs.pl” with parameters “- start -300 -end 300” (we also tested a larger region with “-start -2000 -end 2000” and found similar results).

To identify whether specific motifs were enriched in the promoter regions of differentially expressed genes, we first downloaded the PSWM files for the FOS, PBX3, and PQM-1 binding motifs from Homer^42^. We used the ChIPSeeker^43^ function “getPromoters” with parameters “TxDb=TxDb.Celegans.UCSC.ce11.refGene, upstream=300, downstream=300” to assign promoters to genes from lists of differentially expressed genes generated from the RNA-seq data with a statistical significance threshold of FDR < 0.05 (we also tested a larger region with “upstream=2000, downstream=500” and found similar results). The resulting bed files were converted to FASTA format with bedtools “getfasta”^44^ to be made compatible with the MEME suite AME (McLeay and Bailey, 2010). The MEME Suite AME^45^ (with default parameters “Average odds score” and “Fisher’s exact test” using shuffled input sequences as the control) was used with to determine statistical enrichment of the chosen motifs. We found that the PQM-1 motif was significantly enriched at the male sperm upregulated (*p* = 3.87 × 10^−22^) and downregulated (*p* = 1.23 × 10^−20^) genes, the male seminal fluid upregulated (*p* = 5.87 × 10^−3^) and downregulated (*p* = 9.8 × 10^−19^) genes, and the male pheromone upregulated genes (*p* = 4.93 × 10^−6^).

---

**Extended Data Figure 1:**
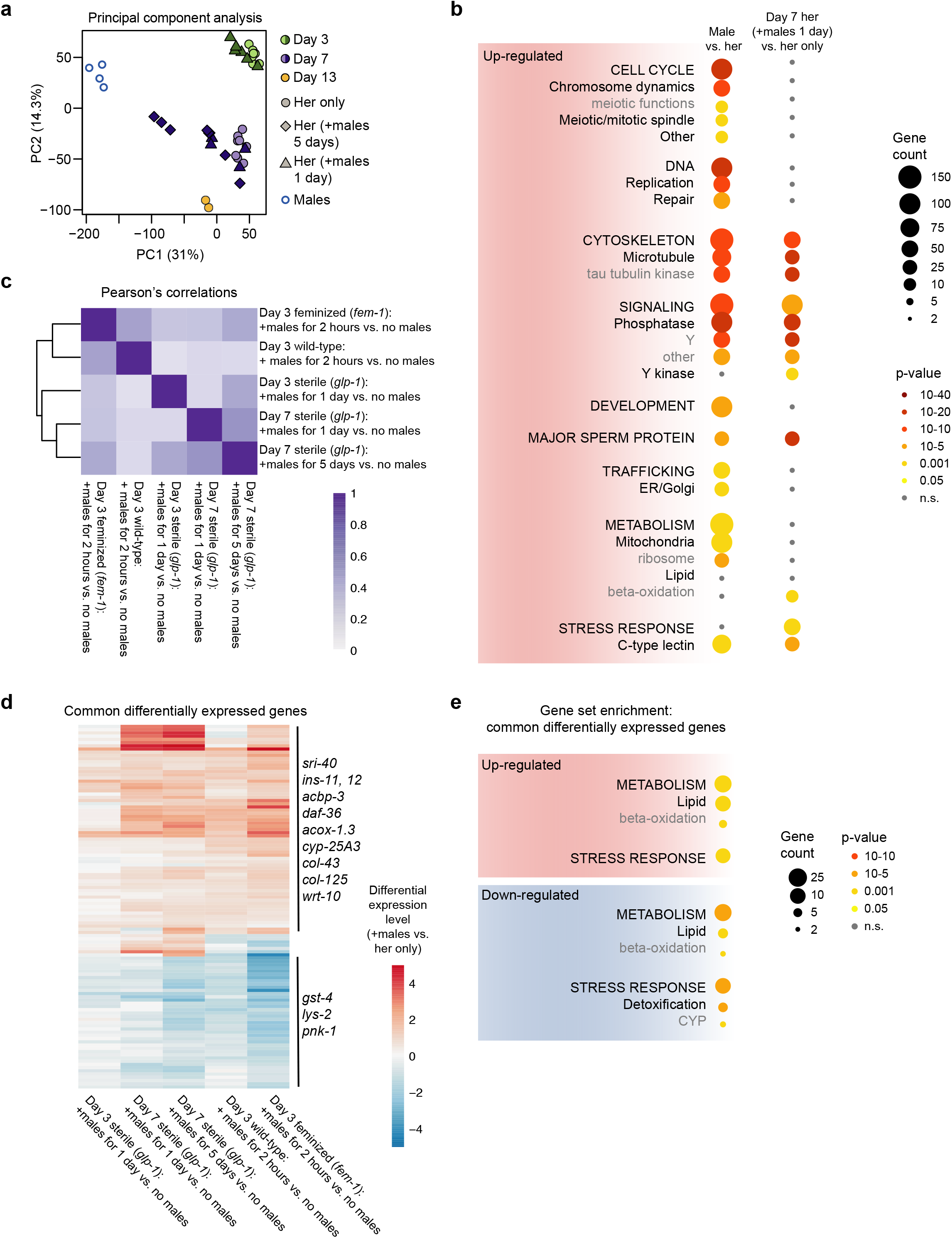
RNA-seq of *C. elegans* hermaphrodites and males. (a) Principal Component Analysis of the normalized read counts of the transcriptomes of *glp-1(e2144)* hermaphrodites and wild-type males prior to removal of the male-enriched genes (see Material and Methods). (b) Enriched gene categories in the genes expressed more highly in wild-type males versus *glp-1* hermaphrodites (left) and the genes expressed more highly in *glp-1* hermaphrodites that experienced a sexual interaction for one day (starting at day 6 of life) versus no sexual interaction (right). The number of differentially expressed genes in each gene set is indicated by the size of the circle and the significance of the enrichment by the color of the circle. Gene annotations are nested with the broadest categories listed in all capital letters and the middle categories listed with the first letter capitalized, and the most specific categories in grey. These two sets of differentially expressed genes share several enriched gene categories, including those linked with sperm such as the major sperm proteins, tau tubulin kinases^33^, and phosphatases/kinases^48^ suggesting that male sperm-derived transcripts are detected by whole-worm RNA-seq of mated hermaphrodites. (c) Pearson’s correlation of the DEseq2 calculated log_2_(fold change) differential expression of all shared, detected genes in wild-type hermaphrodites, sterile *glp-1(e2144)* individuals, and feminized *fem-1(hc17)* individuals following a sexual interaction with males versus no sexual interaction. Data are from this manuscript and Booth *et al. eLife* 2019. (d) Heatmap of the genes that are differentially expressed in at least three of the five conditions. The data are displayed as log_2_(fold change) and the complete results of the differential expression analysis can be found in Supplementary Table 2 and Booth *et al. eLife* 2019. (e) Enriched gene categories in the genes that are upregulated (red) or downregulated (blue) following sexual interactions with males in at least three of the five datasets from *glp-1* sterile hermaphrodites (this manuscript), *fem-1* feminized individuals, and WT hermaphrodites^13^. Complete differential expression and gene set analysis results are in Supplementary Tables 2 to 5.

**Extended Data Figure 2:**
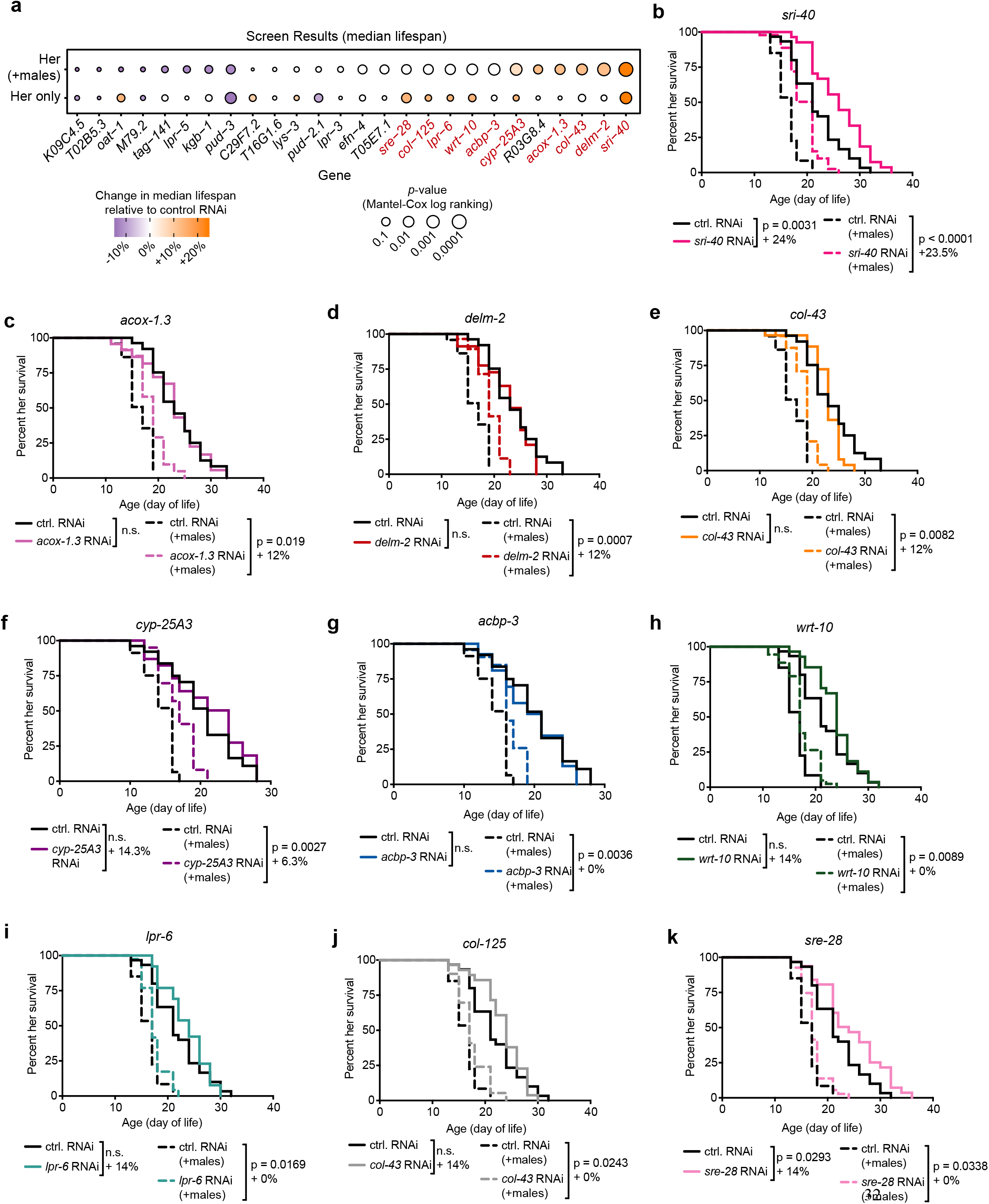
RNAi screen results. (a) A summary of the results of the RNAi-based screen to identify functionally important male-induced gene expression changes. The colors of the circles indicate the change in median lifespan relative to control, empty vector RNAi. The sizes of the circles indicate the *p*-values calculated using Mantel-Cox log ranking. Hits from the screen are shown with red gene name labels. Genes are ordered from smallest to largest change in median lifespan in the presence of males, using *p*-value to break ties. (b-k) The Kaplan-Meier survival curves of the screen hits (*p* ≤ 0.05 when comparing control RNAi versus gene of interest RNAi in the presence of males). Control RNAi treatment is in black and gene of interest RNAi treatment is plotted as colored lines. Hermaphrodite lifespans measured in the presence of males are shown as dashed lines. Several of the control RNAi curves are identical between panels because these genes were tested together in a group of blinded conditions with a single control RNAi condition. For the screen, 23-59 wild-type hermaphrodites were tested. For each lifespan curve comparison, the *p*-value (Mantel-Cox log ranking) and percent change in median lifespan are shown. See Supplementary Table 1 for Supplementary Figure 1. Extended statistics as well as the groupings of the genes tested in the screen.

**Extended Data Figure 3:**
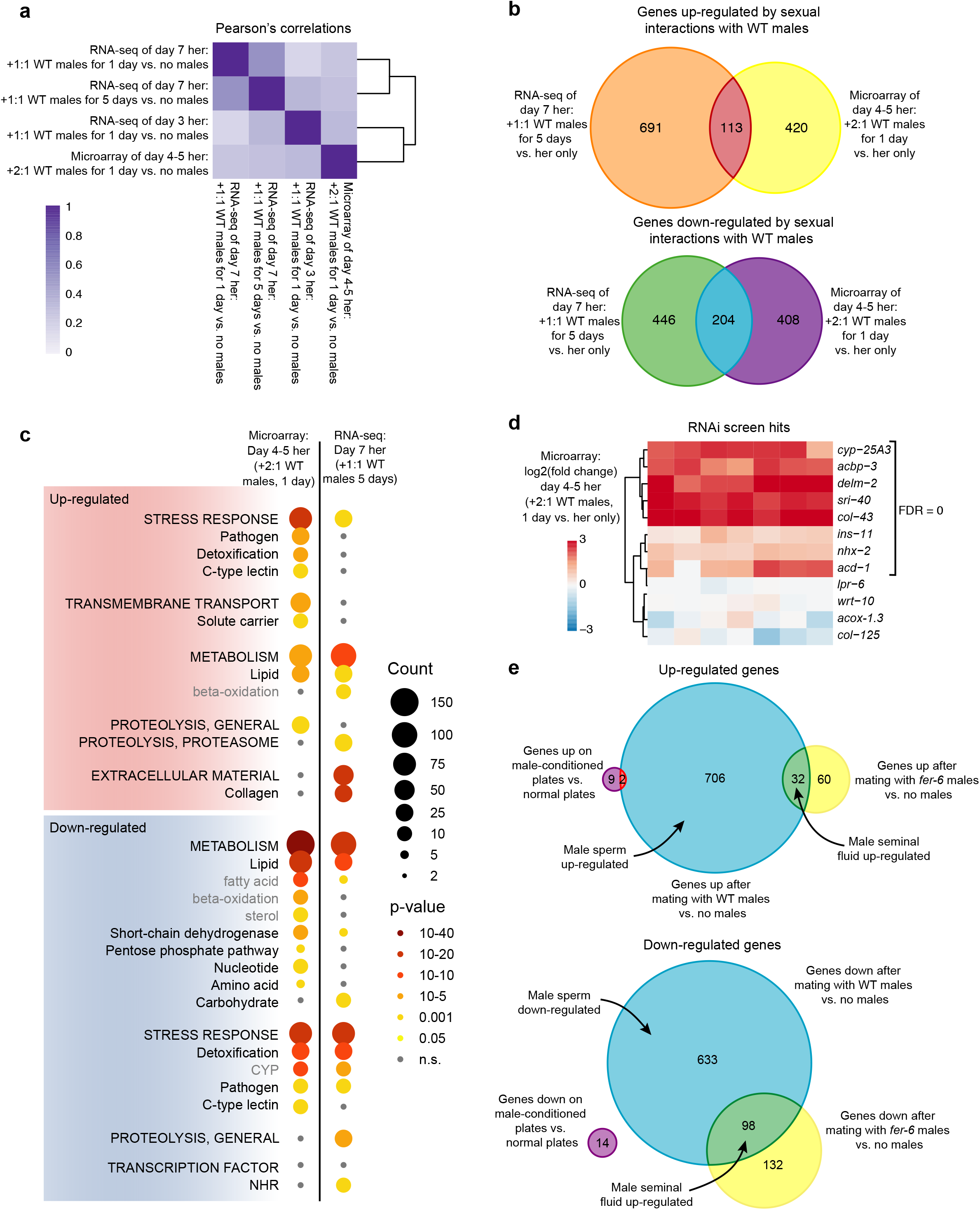
Comparison of RNA-seq and microarray results. The RNA-seq and microarray experiments were performed by two different researchers at different institutes using different experimental setups (*e.g.* the ratio of males to hermaphrodites and the age of the hermaphrodites) and different alleles of *glp-1*. For complete details, see the Materials and Methods. (a) Pearson’s correlation scores comparing the wild-type male-induced fold-change in hermaphrodite gene expression detected in the *glp-1* hermaphrodite RNA-seq datasets and the *glp-1* hermaphrodite microarray dataset. (b) Venn diagrams of the wild-type male-induced differentially expressed hermaphrodite genes detected by microarray and RNA-seq. The number of overlapping differentially expressed genes is higher than expected by chance (upregulated: *p* = 6.15 × 10^−21^, downregulated: *p* = 1.69 × 10^−93^, hypergeometric test). (c) Enriched gene categories in the hermaphrodite genes that are upregulated (red) and downregulated (blue) by wild-type males in the microarray and the RNA-seq results. The number of differentially expressed genes in each gene set is indicated by the size of the circle and the significance of the enrichment by the color of the circle. Gene annotations are nested with the broadest categories listed in all capital letters and the middle categories listed with the first letter capitalized, and the most specific categories in grey. The gene category enrichment for all datasets, including those not shown here, can be found in Supplementary Tables 3 and 9. (d) A heatmap of the microarray results from *glp-1* hermaphrodites mated with WT males for one day. The genes that we identified as functionally important for male-induced demise (Fig. 1i and Supplementary Figure 1. Extended Data Fig. 2) are shown, including *ins-11^1^*. The majority of genes that were differentially expressed in our RNA-seq and identified as functionally important in our screen were also differentially expressed in the microarray experiment. The four genes that were not significantly up-regulated under the different experimental conditions used for the microarray were not investigated further. *sre-28* was not detected in the microarray. (e) Venn diagrams of the significantly up- and down-regulated hermaphrodite genes detected by microarray following one day of interacting (mating) with WT males or *fer-6(hc6)* sperm-less males (blue and yellow, respectively) or five days on male-conditioned plates (i.e. male secreted compounds and pheromones, purple). A complete list of all differentially expressed genes from the RNA-seq and microarrays can be found in Supplementary Tables 2, 6, 7, and 8.

**Extended Data Figure 4:**
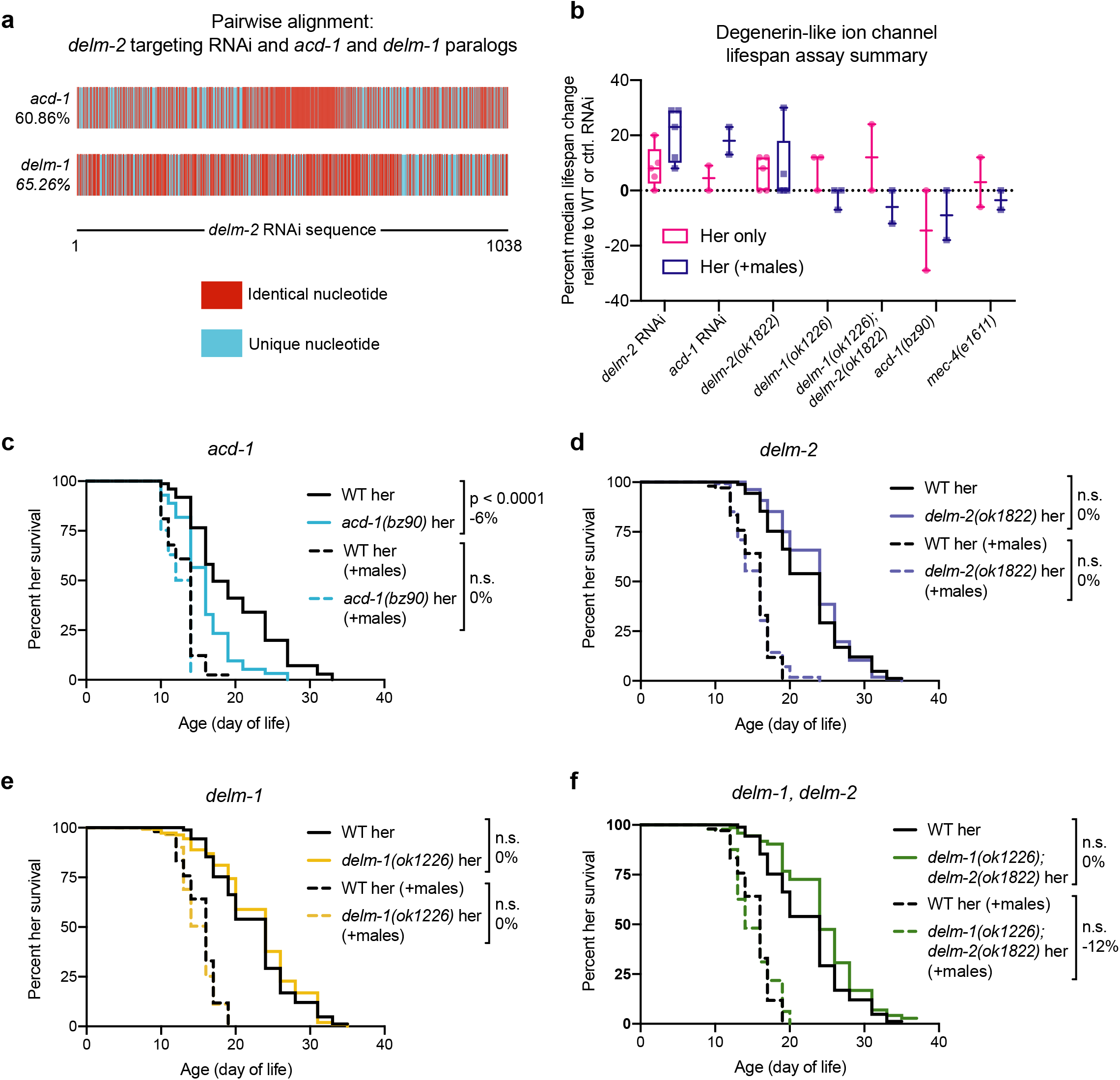
Degenerin-like ion channel lifespan assays. (a) DNA sequence alignment of the *delm-2* RNAi targeting sequence from the Ahringer RNAi library^35^ with the aligning portions of the *delm-1* and *acd-1* unspliced transcripts. (b) A summary of the male-induced demise lifespan assay results testing the role of the degenerin-like ion channel paralogs *acd-1*, *delm-1*, and *delm-2*. Lifespan data (median lifespan change relative to control) for each biological replicate are shown as individual data points and summarized with a box plot. Box plot whiskers show minimum and maximum and the line in the box indicates the median of the biological replicates. (c-f) Loss of function mutations in *acd-1, delm-1*, or *delm-2* alone (panels c-e) were not sufficient to extend hermaphrodite lifespan in the presence or absence of males, nor did *delm-1, delm-2* double mutation (panel f). We note that RNAi targeting either *delm-2* or *acd-1* is sufficient to extend hermaphrodite lifespan (Fig. 2e and Fig. 4d, g). 89-138 hermaphrodites were tested in panels b-f. Lifespans are plotted as Kaplan Meier survival curves and *p*-values calculated by Mantel-Cox log ranking. Percent change in median lifespan compared to control is shown in panels c-f. See Supplementary Table 1 for Supplementary Figure 1. Extended statistics.

**Extended Data Figure 5:**
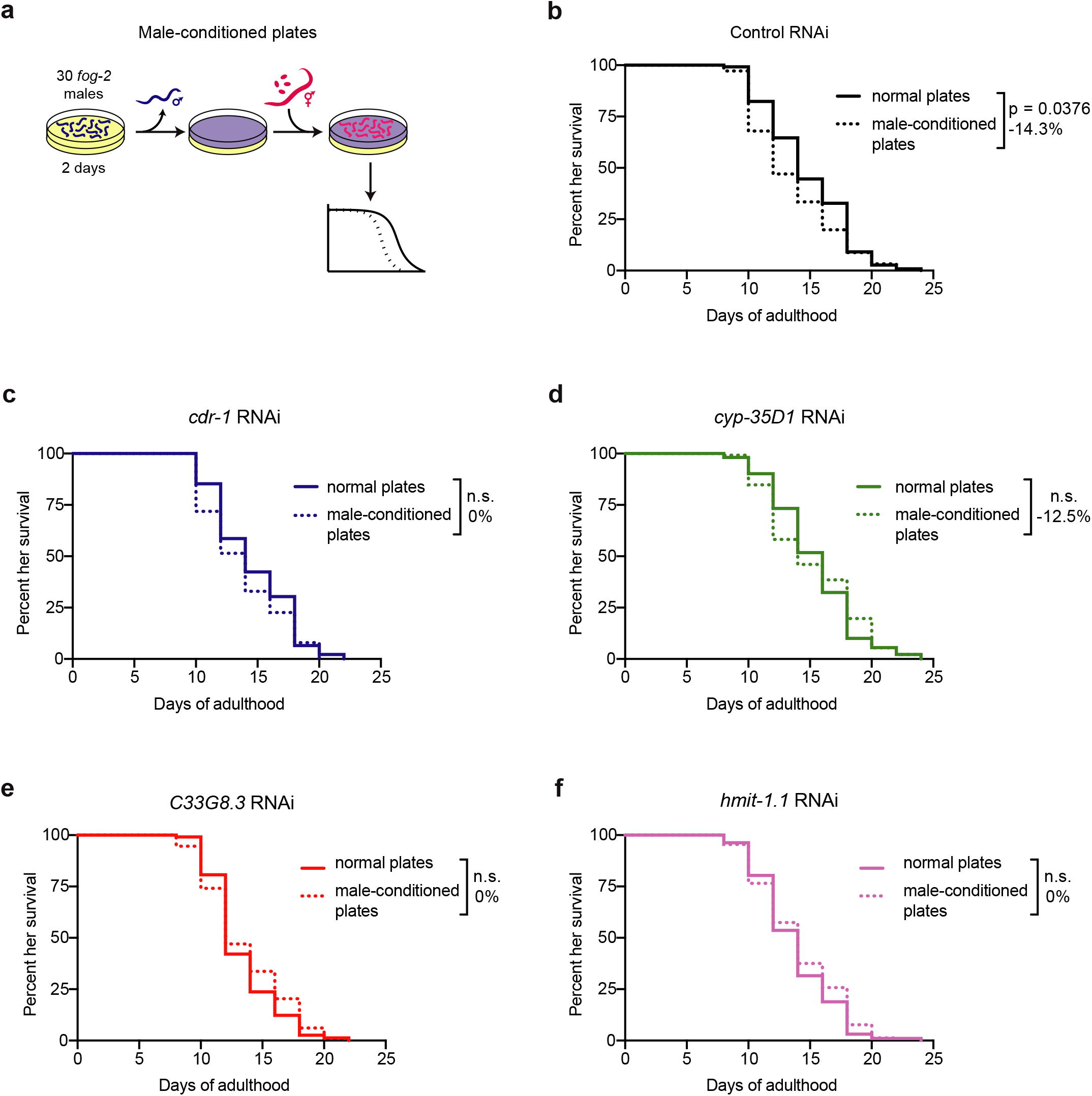
Male-conditioned plate screen results. (a) A scheme describing the experimental set up for the male-conditioned plate lifespan assays. For complete details, see the Materials and Methods. (b-f) The lifespans of hermaphrodites fed control empty vector RNAi (b) or RNAi against the male-conditioned plate induced genes *cdr-1* (c), *cyp-35D1* (d), *C33G8.3* (e), or *hmit-1.1* (f). Hermaphrodite worms were either exposed to media conditioned by 30 *fog-2(q71)* males for 2 days (removed prior to placing the hermaphrodites on the plate) (dotted lines) or were placed on normal plates (solid lines). Results are plotted as Kaplan-Meier survival curves and *p*-values calculated using Mantel-Cox log ranking. Percent change in median lifespan compared to normal plates is shown in each panel. 118-132 hermaphrodites were tested in each condition. See Supplementary Table 1 for a complete list of all lifespan assay results.

**Extended Data Figure 6:**
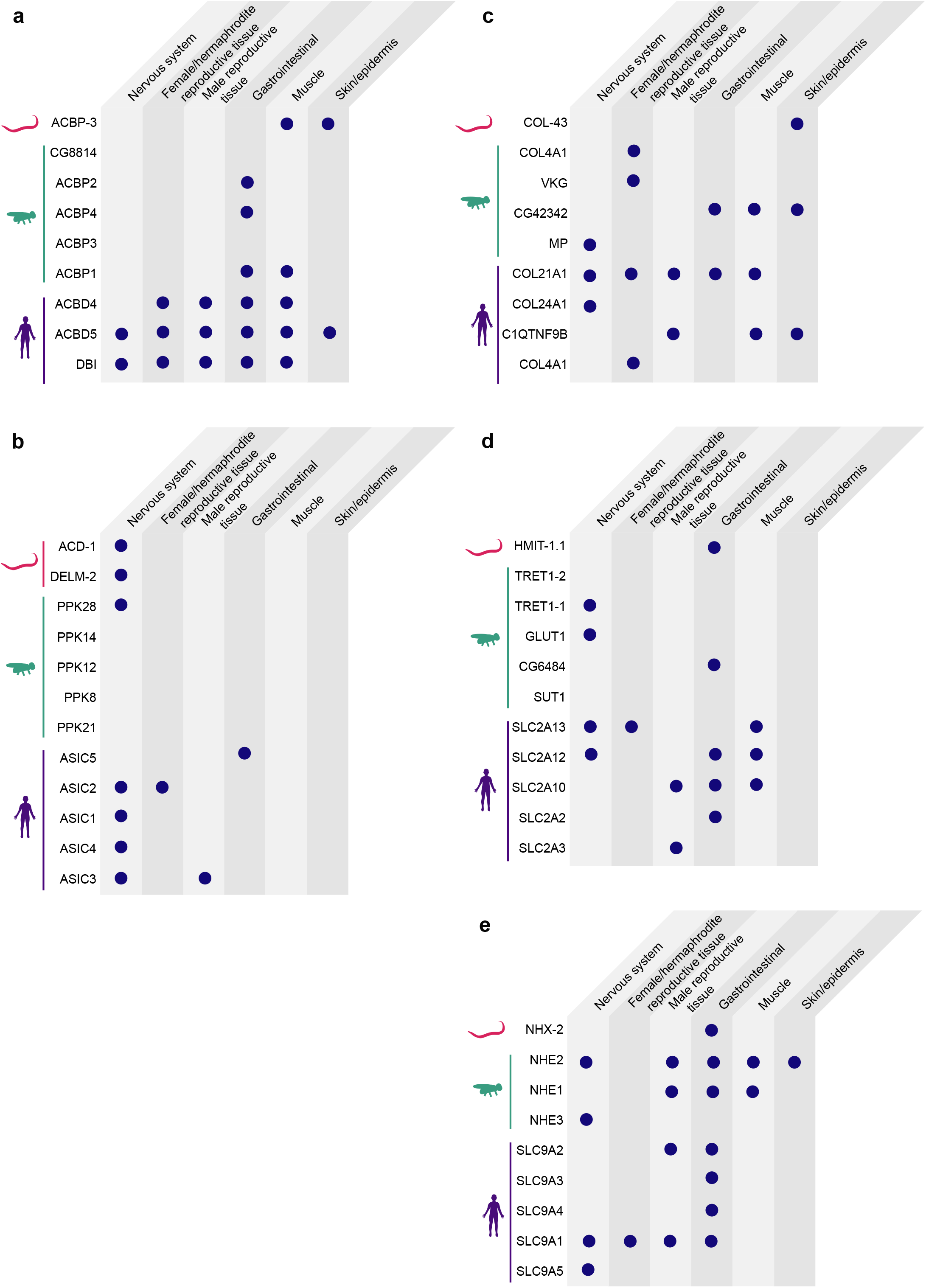
Conservation of male-induced demise genes. The top five BLASTp hits in *Drosophila melanogaster* and humans for the *C. elegans* male-induced demise genes *acbp-3* (a), *acd-1* and *delm-2* (b), *col-43* (c), *hmit-1.1* (d), and *nhx-2* (e). The tissue-specific protein expression is indicated for each ortholog in *C. elegans* (www.wormbase.org, version WS275), *Drosophila* (www.flybase.org, version 2020_01), and humans (www.proteinatlas.com^46,47^). A portion of these data are also presented in Figure 2g-i.

**Extended Data Figure 7:**
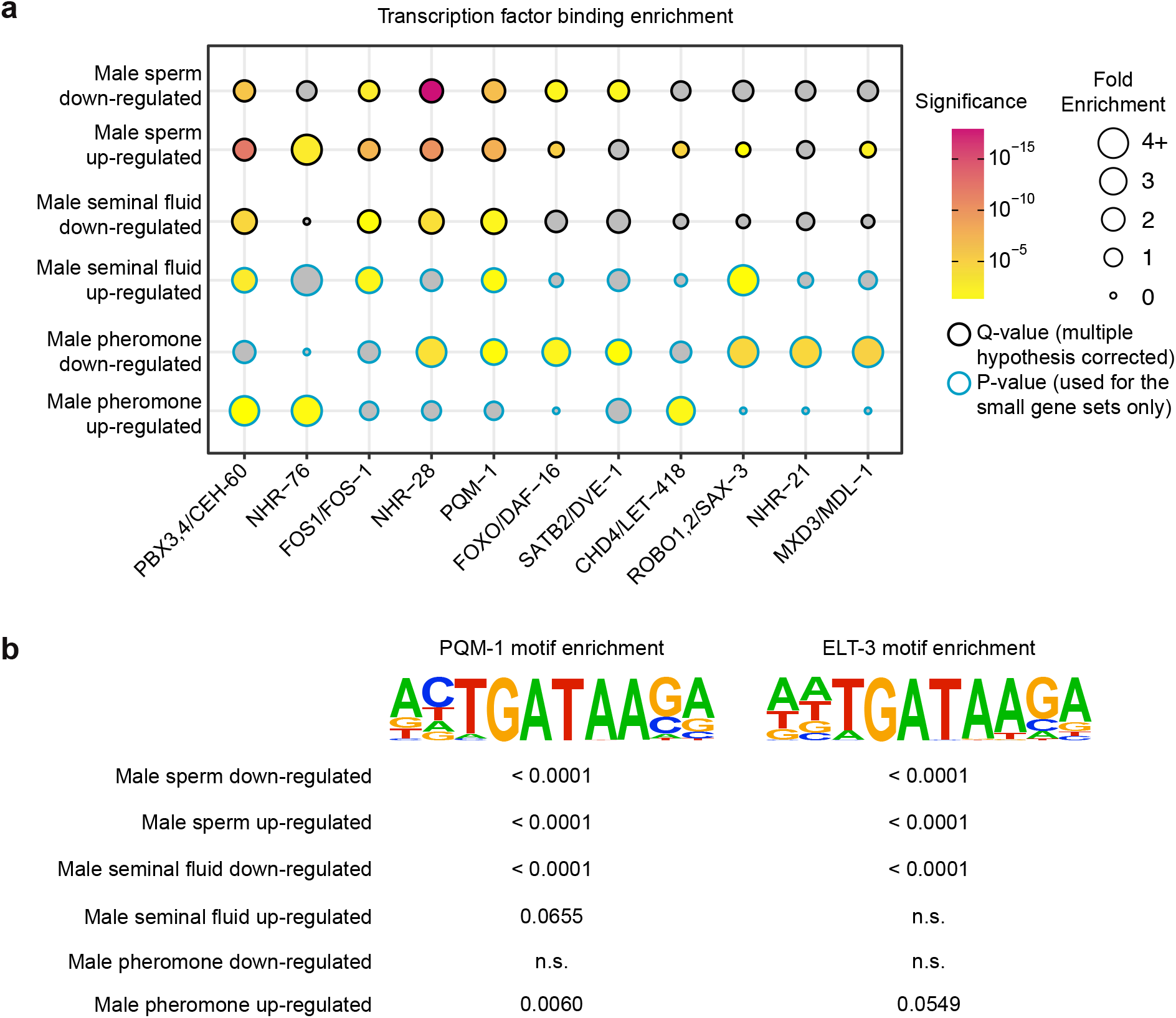
Transcription Factor Binding Enrichment. (a) The enrichments of selected transcription factor binding peaks within 2kb +/− the transcription start site of the male sperm, seminal fluid, and pheromone downregulated and upregulated genes are shown. Fold enrichment at the gene set of interest compared to genome-wide is indicated by the size of each circle and Q-value (false discovery corrected *p*-value) or *p*-value by the color of each circle. A complete list of the transcription factor binding enrichment is presented in Supplementary Table 10. (b) Enrichment of “known” transcription factor binding motifs within 300bp +/− the transcription start site of the differentially expressed genes. The Bonferroni-corrected *p*-values for the motif enrichment for each gene set is shown.

**Extended Data Figure 8:**
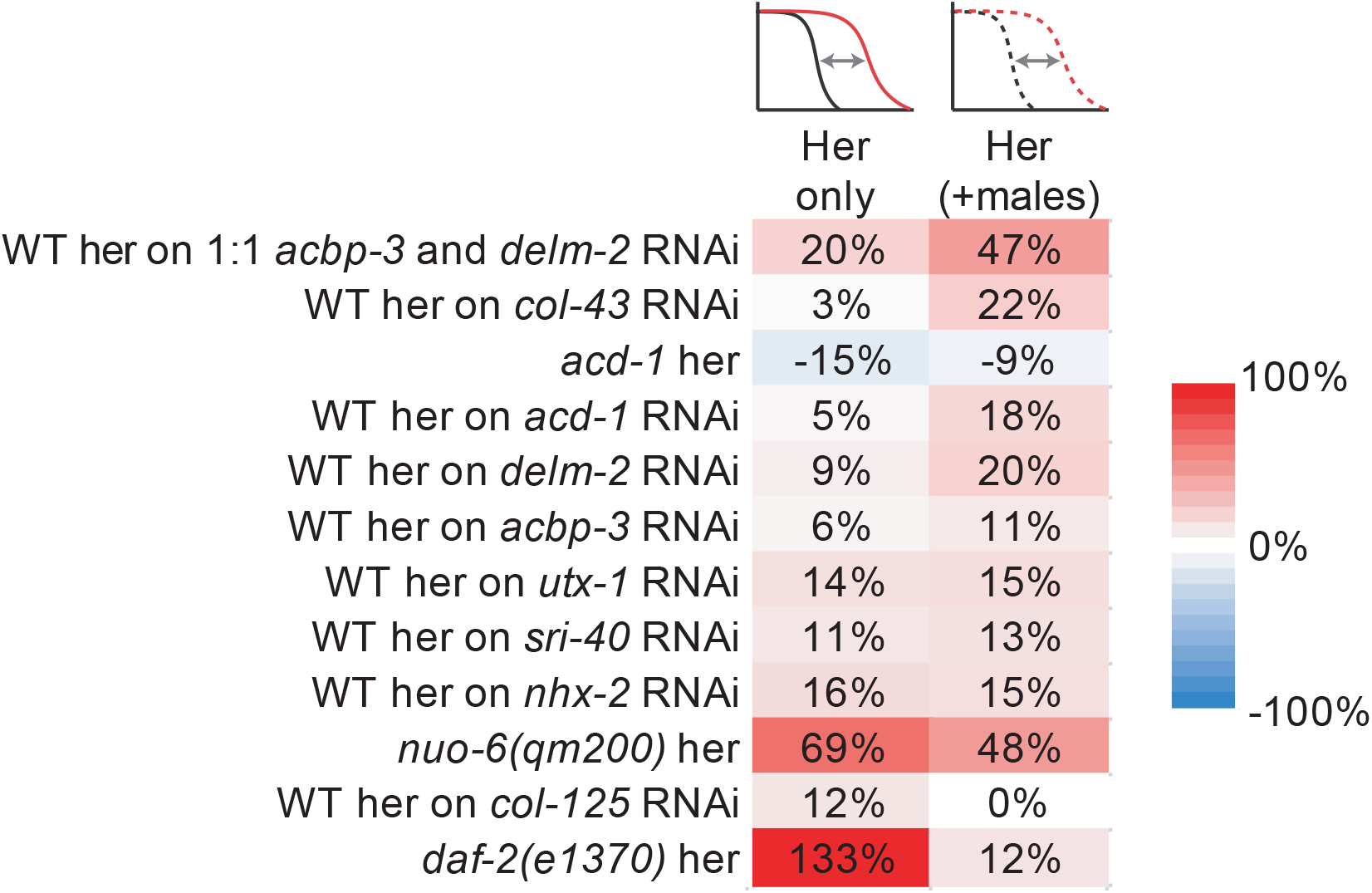
Male-induced demise and longevity. A summary of the percent loss of lifespan in the presence of males in the mutant or RNAi knock-down compared to wild-type. Positive values indicate that the genetic manipulation resulted in greater protection from male-induced demise whereas negative values indicate hypersensitivity to male-induced demise. The *daf-2(e1370)* results were published previously^1^. All other lifespan results are from this manuscript (see Supplementary Table 1 and Supplementary Table 11).

## Supplementary Tables

**Supplementary Table 1: Lifespan assay results**

**Supplementary Table 2: DESeq2 calculated RNA-seq differential expression after removal of male-enriched transcripts**

**Supplementary Table 3: Gene set enrichment of the RNA-seq differentially expressed genes (after removal of male-enriched transcripts)**

**Supplementary Table 4: DESeq2 calculated RNA-seq differential expression**

**Supplementary Table 5: Gene set enrichment of the RNA-seq differentially expressed genes**

**Supplementary Table 6: Microarray SAM results for WT male-mated hermaphrodites**

**Supplementary Table 7: Microarray SAM results for spermless (*fer-6)* male-mated hermaphrodites**

**Supplementary Table 8: Microarray SAM results for male-conditioned plate exposed hermaphrodites**

**Supplementary Table 9: Gene set enrichment of the microarray differentially expressed genes**

**Supplementary Table 10: Transcription factor ChIP-seq enrichment**

**Supplementary Table 11: Comparison of lifespans in the presence and absence of males**

## Notes

### Competing Interest Statement

The authors have declared no competing interest.

